# Preparation and application of YFV-17D-derived Anterograde Trans-neuronal Viral Vectors for Neuroscience Research

**DOI:** 10.64898/2026.01.08.698485

**Authors:** Tanvi Panchumarthy, Lizbeth Munoz, Uzair Saleem, Ai-Vy Le, Ian Nelson, Ying Li, Xiangmin Xu, Wei Xu

## Abstract

Trans-neuronal viruses that spread between synaptically connected neurons have become invaluable tools in neuroscience, enabling circuit mapping and targeted delivery of genetic material to cells within defined pathways. Although most available trans-neuronal viral tracers propagate retrogradely, an anterograde trans-neuronal virus that spread from presynaptic to postsynaptic neurons would substantially expand experimental capabilities. We previously demonstrated that YFV-17D—a live attenuated yellow fever vaccine used clinically for decades—spreads anterogradely along neuronal circuits. We further developed a suite of recombinant YFV-17D–based vectors for diverse applications, including tracing monosynaptic projectome of defined neuronal cell types, multiplex mapping of parallel pathways, and trans-neuronal genetic manipulation with minimal neurotoxicity. We have now tested and optimized procedures for vector production and application. Based on these optimizations, here, we present a two-part guide for preparing and deploying this viral vector system to map brain connectivity. The first part describes the molecular biology and cell culture workflows required to generate high-quality viral vectors, yielding titers of ∼1 × 10^10 to 5 × 10^11 genomic copies per milliliter within approximately two weeks. The second part outlines optimized animal procedures, including intracranial injections, perioperative care, and experimental considerations tailored to specific aims and vector variants. Depending on the application, efficient trans-neuronal tracing can be achieved within 1–4 weeks following delivery.

## Introduction

Viral vectors are indispensable tools for both neuroscience research and clinical applications. Trans-neuronal viral vectors are particularly valuable for mapping neuronal connectivity and functionally manipulating neuronal circuits^1–4^. Anterograde trans-neuronal viruses, for example, can be used to elucidate the “projectome“—the direct downstream neurons throughout the brain of a specific neuronal population. They can also selectively label neurons directly innervated by a targeted neuronal group, without affecting neighboring neurons within the same brain region. However, the availability of trans-neuronal viruses capable of crossing synapses in the anterograde direction remains limited. Recently we discovered that YFV-17D, the well-established yellow fever vaccine, is a promising candidate for anterograde trans-neuronal tools^5^. It has several advantageous features: (1) As one of the most widely used vaccines^6^, it is safe for laboratory handling; (2) it efficiently crosses synapses in the anterograde direction; (3) it has broad neuronal tropism; (4) it can be modified easily due to its compact genome (11kb) ^7^; and (5) it demonstrates lower neurotoxicity compared to many replication-capable viruses. To enhance the functionality of the original replication-capable YFV-17D, we constructed several derivatives using a trans-complementation strategy^8,9^ (Fig.1). In the YFV^ΔNS1^ system, we targeted the viral replication process. Inducible NS1 expression via helper AAVs enables selective anterograde tracing with minimal neurotoxicity, allowing for genetic manipulation of postsynaptic neurons. In another system, YFV^ΔCME^, we leveraged viral particle assembly to enable monosynaptic tracing, in which the virus reaches only directly connected neurons, without spreading farther. This system is particularly suitable for mapping monosynaptic projections throughout the brain. Since the initial use of YFV-17D in neuronal circuit studies, we have systematically optimized protocols for virus packaging, purification, and in vivo application in animal models to achieve high efficiency and reproducibility. This paper describes these optimized protocols.

**Fig. 1.**
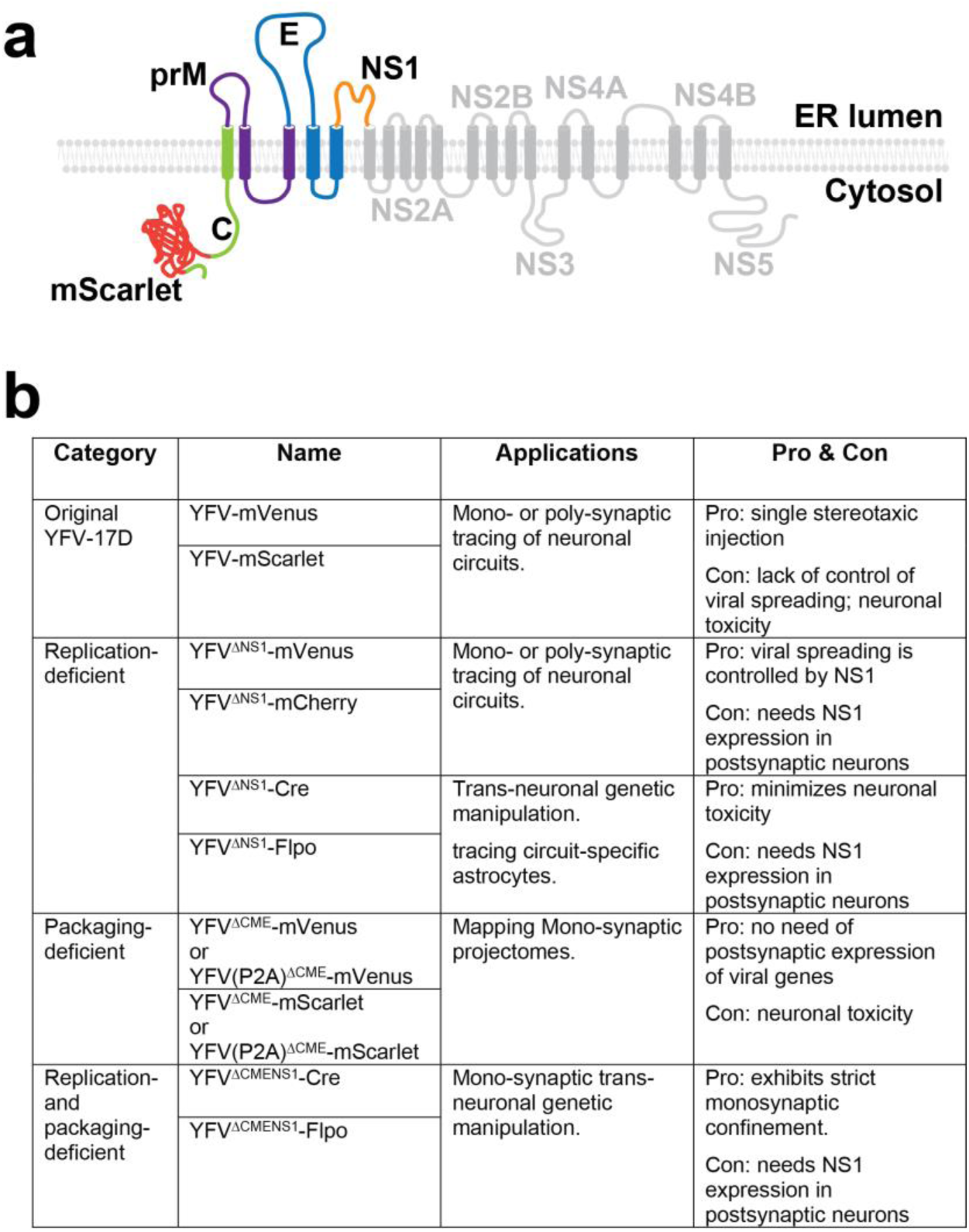
Selection of Viral Vectors. (**a**) Genes Encoded by the YFV-17D Genomic RNA. The YFV-17D genome contains a single open reading frame (ORF) that translates into a polyprotein on ER membrane, which is subsequently cleaved into ten distinct proteins. Among these proteins, C, prM, and E are structural proteins assembled into mature virions. In contrast, the nonstructural proteins (NS1 to NS5) are essential for viral replication, assembly, and egress but are not incorporated into virions. Cargo gene(s) can be inserted into the coding sequence of C. Removal of the NS1 gene from YFV-17D (generating the YFV^ΔNS1^ series), rendering the virus incapable of replication unless NS1 protein is supplied *in trans* within the same cells. Removal of the C, prM, and E genes (generating the YFV^ΔCME^ series), preventing the assembly and egress of virions. These functions can be restored only if the C-prM-E proteins are provided *in trans* within the same cells. (**b**) Applications and Limitations of YFV-17D-Derived Viral Vector Versions. Various YFV-17D-derived vectors have been developed, each tailored for specific research applications.

The viral tools derived from YFV-17D can be applied alone or combined with other modern neuroscience techniques to reveal both structural features of the brain—such as neuronal circuit connectivity—and its functional organization, including the roles specific neurons play in brain function. Potential applications include, but are not limited to, the following: (1) Mono- and poly-synaptic tracing of neuronal circuits: YFV-17D carrying fluorescent protein genes can effectively trace neuronal pathways. Typically, it takes about three days for the virus to traverse one order of synaptic connections in the mouse brain, allowing for either mono-synaptic or poly-synaptic tracing by adjusting post-injection timing of perfusion. The expression of fluorescent proteins by YFV-17D can be readily detected without the need for amplification via immunohistochemistry^5^. (2) Mono-synaptic tracing from defined neuronal cell types: Tracing with YFV^ΔNS1^ or YFV^ΔCME^ can be selectively initiated from specific neuronal cell types. These viruses require a trans-complementary expression of the NS1 or CME genes to spread. These genes can be conditionally expressed in specific cell types allowing replication only in these cells, enabling cell-type-specific tracing^5^. (3) Trans-neuronal genetic or functional manipulation of postsynaptic neurons: YFV-17D viruses can be engineered to carry DNA recombinases (like Cre or Flp) or other genetic modifiers that alter gene expression in postsynaptic neurons. Once genetic modification is achieved, viral replication can be stopped to minimize potential neurotoxicity, allowing the study of neuronal function and connectivity in behaving animals^5^. (4) Mapping convergence and divergence in neuronal circuits: By incorporating multiple fluorescent proteins into YFV-17D vectors, researchers can visualize circuit convergence and divergence within the same animal. YFV-17D also holds promise for barcode-based tracing from individual neurons, providing even more detailed insights into circuit convergence and divergence^5,10^. (5) Tracing glial cells associated with specific neuronal circuits: Astrocytic processes play an essential role in tripartite synapses and neural communication. YFV^ΔNS1^, carrying DNA recombinases, can trace astrocytes associated with specific neuronal circuits, enabling circuit-specific astrocyte mapping across the whole brain^11^. (6) Studying the transport of pathogens and spreading of endogenous pathologies. Many infectious pathogens or pathological alterations in many neurodegenerative disorders spread in the brain along brain circuits^12^. With their well-defined features for studying neuronal circuits, YFV-17D vectors can be used to study the molecular and cellular machinery involved in pathogenic progression and reveal the defects underlying those disorders.

## Results

Our optimization of the YFV-17D-based system has been focusing on the following three aspects: **(1) Expanding the Viral Vector Toolkit**. In addition to the original YFV-17D, we have developed multiple YFV-17D-derived vector variants with trans-complementation strategy^9^, each tailored for specific applications. Researchers should carefully evaluate the advantages and limitations of each vector to select the most suitable one for their experimental objectives (Fig. 1). Some experimental needs may extend beyond the capabilities of existing vectors, such as incorporating new cargo genes or adding tags and barcodes. In such cases, standard molecular biology techniques, including restriction digestion and ligation or Gibson assembly, can be used to modify these vectors. Several restriction sites incorporated into the vectors facilitate molecular cloning (Fig. 2). In earlier versions of the viral vector, an F2A (Foot-and-mouth disease virus 2A peptide) sequence and an approximately 220 bp sequence derived from *Uba52* were inserted between the cargo gene and the viral gene C to achieve bicistronic gene expression. Recently, we tested some updated viral vectors in which these components were replaced with a single P2A (Porcine teschovirus-1 2A peptide) sequence. Compared to the earlier YFV^ΔCME^-mScarlet vector, the P2A-based version, YFV(P2A)^ΔCME^-mScarlet, exhibited similar or slightly higher titers and trans-neuronal spreading efficiency. Additionally, this modification freed up approximately 200 bp of space, allowing for the inclusion of larger cargo genes. The region between the 5’ UTR and the cargo gene contains an 81-bp sequence essential for viral RNA replication and cyclization. When modifying viral vectors derived from YFV-17D, this region must remain intact.

**Fig. 2.**
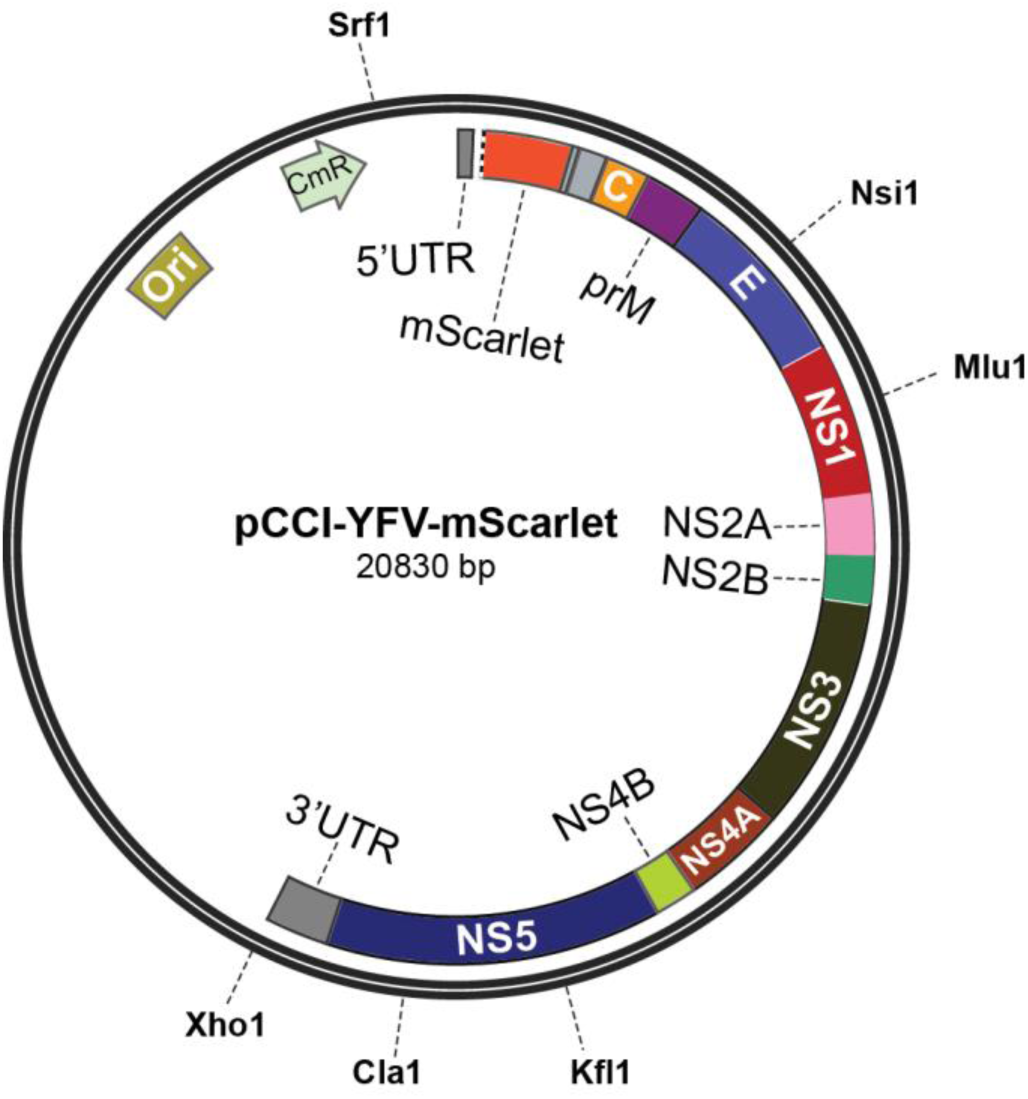
Map of the pCCI-YFV-mScarlet Plasmid, an Example of YFV-17D-Derived Viral Vectors. The restriction enzymes indicated on the map each have a single-cut restriction site, can be used for further modifications of the vector as needed. The XhoI site is located immediately downstream of the YFV-17D genomic sequence and can be used to linearize the plasmid for *in vitro* transcription. Note that the region between the cargo gene (mScarlet) and viral gene C contains an F2A sequence and a segment derived from *Uba52* to enable bicistronic expression. Our recent tests have demonstrated that these two components can be replaced with a single P2A sequence, achieving similar viral titers and trans-neuronal spreading efficiency. Abbreviations: Ori: Origin of replication; CmR: Chloramphenicol resistance gene; UTR: Untranslated region.

**(2) Optimizing generation methods for YFV-17D viruses and helper AAVs.** (Fig.3 and Fig.4). YFV-17D is an enveloped RNA virus, whose genomic RNAs replicate in cultured cells before assembling with structural proteins (C-prM-E) into virions, which are then released into the culture medium. The preparation workflow (Fig. 3) begins with plasmid DNA amplification, followed by in vitro transcription of the plasmid to generate genomic RNAs of YFV-17D or its variants. These RNAs are then transfected into cell lines to initiate viral replication and egress. The final step involves concentrating or purifying the virus from the culture medium. Key considerations during this process include the scale and purity of viral preparations. Handling YFV-17D viral vectors and associated reagents must adhere to BSL-2 safety protocols. The final viral titer can be quantified via real-time PCR or cell infection assays. Helper AAVs, which provide critical support for some YFV-17D-derived vectors, can be produced using established protocols. The desired titer (approximately 1E13 copies/ml) and purity should guide AAV preparation efforts. As these methods are widely available^13^, detailed discussions are omitted here.

**Fig. 3.**
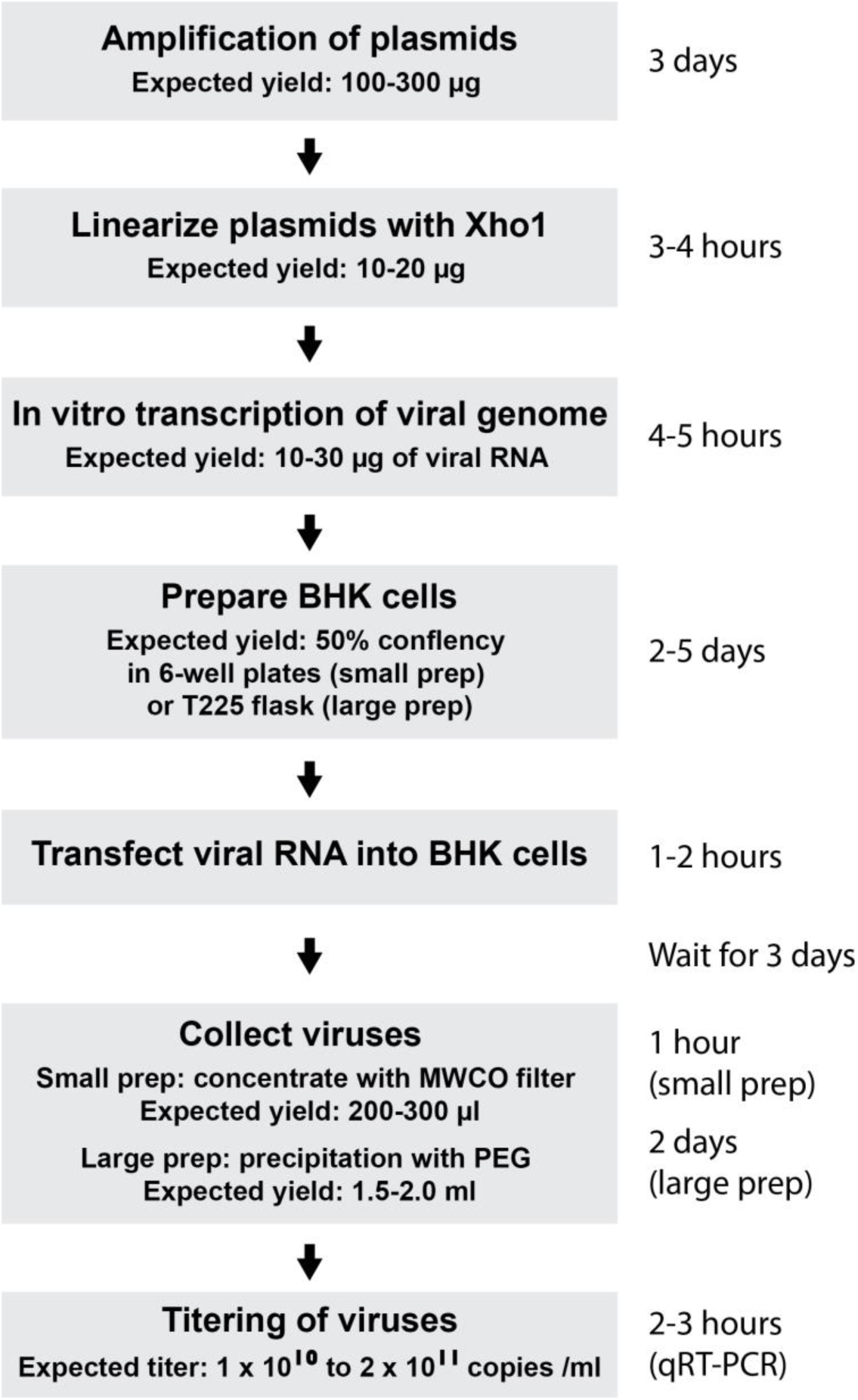
Workflow for Packaging YFV-17D-derived Viral Vectors.

**Fig. 4.**
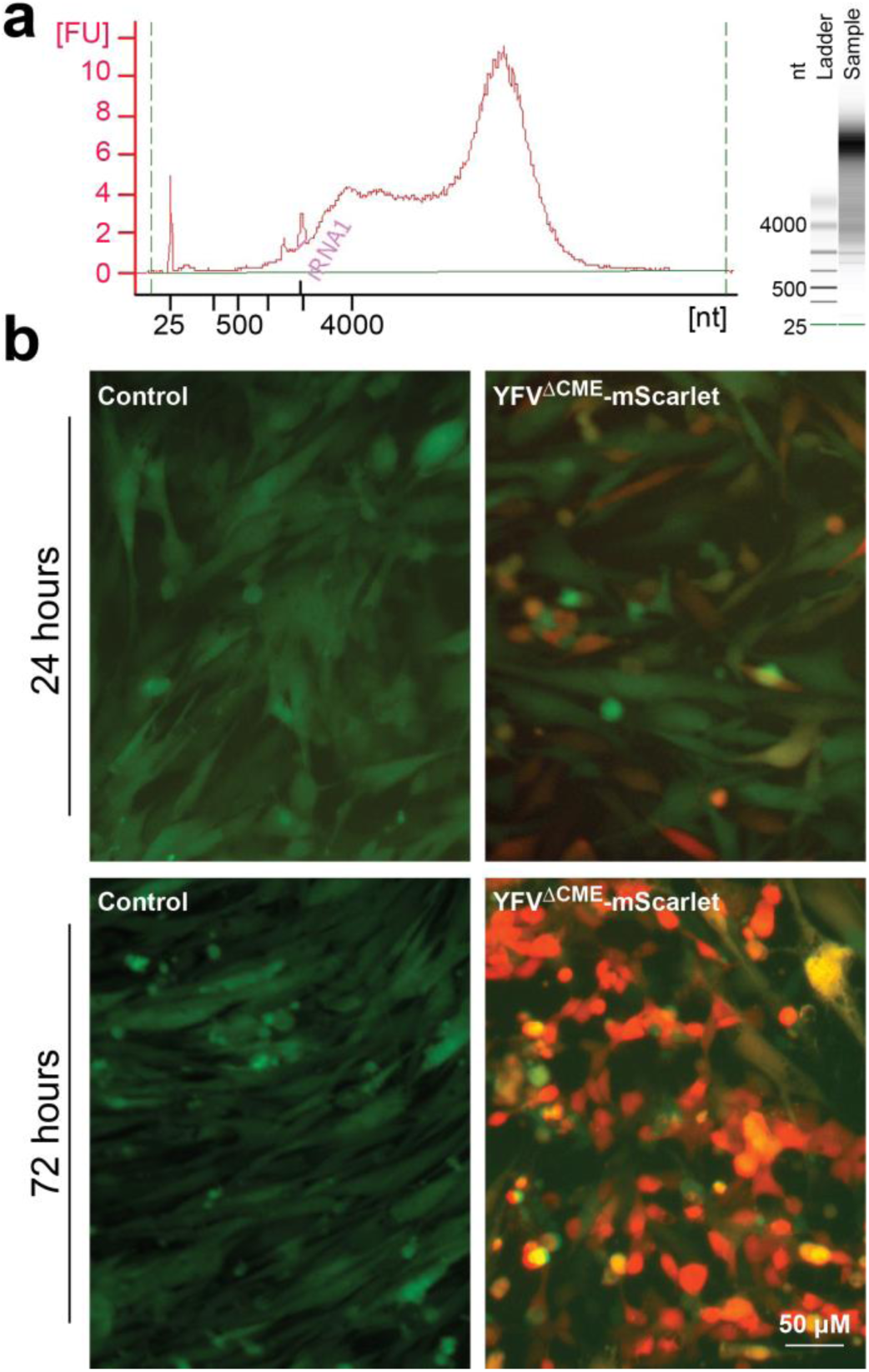
Packaging of YFV-17D-derived viral vectors with BHK cells. (**a**) Example of viral RNA produced via *in vitro* transcription, analyzed with the Agilent Bioanalyzer 2100. (**b**) Transfection of BHK cells stably expressing C-prM-E-NS1-IRES-EGFP (green signal) with viral RNA of YFV^ΔCME^-mScarlet (red signal). Control groups were not transfected with viral RNA. mScarlet fluorescence became detectable 8–9 hours post-transfection and intensified by 24 hours. By 72 hours, the BHK cells exhibited signs of cytopathy (Lost their elongated or star-shaped morphology, becoming rounded and sometimes detaching from the culture flask.)

**(3) Refining the protocol for in vivo administration of YFV-17D-derived vectors.** (Fig. 5 and Fig.6). The application protocols for YFV-17D-derived vectors vary depending on the vector. Key considerations include achieving efficient trans-neuronal spreading and minimizing cellular toxicity. Experiments using the original, replication-capable YFV-17D only need a single intracranial injection. The virus requires 2–3 days to produce detectableexpression of its carried reporter genes (e.g., mVenus or mScarlet) at the injection site. It then takes an additional 2–4 days to spread to the first order connected downstream neurons. The time required for further propagation across each synaptic connection varies depending on the length of the axonal projections and other biological features of the neurons. For most genetically modified YFV-17D vectors (excluding the original replication-capable YFV-17D), the procedure typically involves two sequential stereotaxic intracranial injections. The first injection delivers helper AAVs to express genes required for YFV-17D replication or packaging. The second injection introduces the modified YFV-17D vector. The optimal interval between these injections depends on the specific vector and has been empirically determined (summarized in Fig. 4). Once transneuronal spreading is achieved, animals can proceed to the planned experiments, which may include anatomical analysis (e.g., brain fixation) or functional analysis of neuronal activity in behaving animals.

**Fig. 5.**
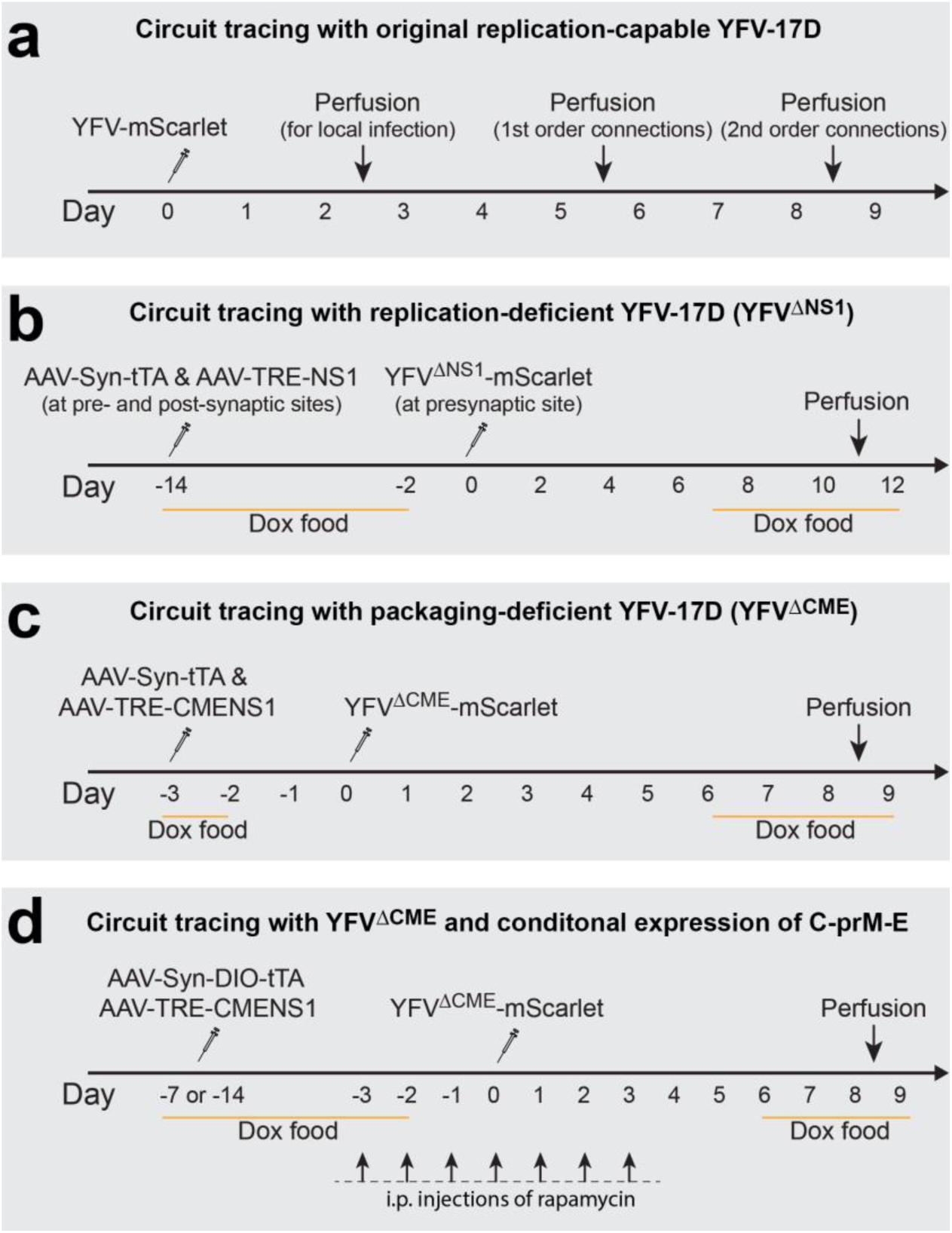
Validated Experimental Procedures for Different Versions of YFV-17D-Derived Viral Vectors. Although these procedures were primarily for neuroanatomy study (involving perfusion and fixation of the brains), the similar time course applies to survival experiments (for example, using YFV^ΔNS1^-Cre to control gene expression in postsynaptic neurons). The “perfusion” dates are approximate and may be adjusted depending on the specific circuits being studied.

**Fig. 6.**
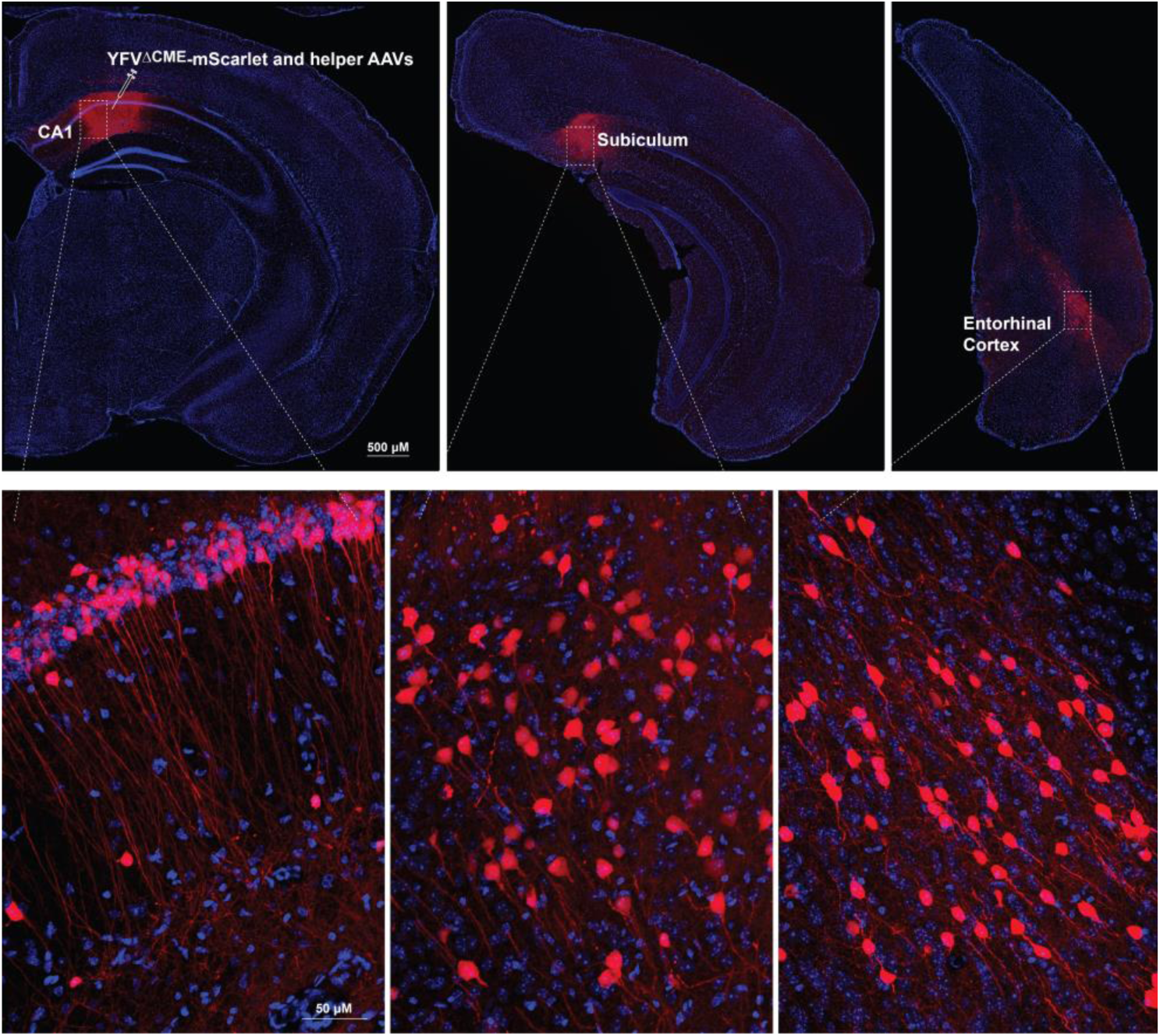
Tracing hippocampal CA1 projection with YFV^ΔCME^-mScarlet. Mouse was injected with YFV^ΔCME^-mScarlet along with helper AAVs (AAV-syn-tTA and AAV-TRE-CMENS1), following the procedure outlined in Fig. 4c. mScarlet-positive neurons were observed at the injection site -- CA1 of the hippocampus, and its downstream brain regions including the subiculum and entorhinal cortex.

## Discussion

Several retrograde trans-neuronal viruses, including rabies virus^1,2,14,15^, pseudorabies virus (PRV) ^16–18^, and vesicular stomatitis virus (VSV) ^19^, have been adapted for neuroscience research, and modified for broader applications. Most retrograde viruses have complementary applications as that of YFV-17D. Previously, the H129 strain of herpes simplex virus (HSV) was the best-characterized anterograde virus^20–23^. Recently, adeno-associated viruses (AAV1 and AAV9) were found to spread across synapses in an anterograde manner^24,25^. Comparatively, the engineered YFV-17D systems offer several advantages: (1) Anterograde directionality: YFV-17D shows no axonal uptake and spreads primarily in the anterograde direction within the first 8–9 days post-injection, with delayed retrograde transport appearing only after 10–15 days. Inducible replication of YFV^ΔNS1^ eliminates this delayed retrograde transport. However, H129 and the aforementioned AAVs exhibit axonal uptake and retrograde transport^26–29^. (2) Synaptic specificity: several lines of evidence show the synaptic specificity of YFV-17D in its trans-neuronal transport, enhancing its precision for circuit mapping^5^. (3) Diverse applications: the YFV-17D vectors can be used for either mono-synaptic or controlled, stepwise poly-synaptic tracing. (4) Minimal neuronal toxicity: constraining YFV-17D replication to a limited time window helps avoid neuronal toxicity, allowing traced neurons to be used for whole-cell patch-clamp recordings or chronic calcium imaging. (5) Targeting astrocytes: YFV-17D can trace astrocytes associated with specific circuits at the whole-brain level, which makes it ideal to study the circuit-specific roles of astrocytes^11,30–33^. (6) Safety: YFV-17D presents minimal biosafety concerns as a well-established clinical vaccine. The generation of the replication- or packaging-deficit versions further reduces these concerns^6,34^.

While the new viral system derived from YFV-17D offers several advantages, it also presents a few limitations that warrant further improvements: (1) **Significant neuronal toxicity in replication-capable versions**. Although neuronal toxicity can be mitigated by using replication-deficient YFV-17D viral vectors, the replication-capable versions still demonstrate significant neuronal toxicity 12–14 days post-injection into the brain. Consequently, experiments using these versions must be carefully timed. (2) **Cell-type-specific tracing may be confounded by local infection**. Although the spreading of either YFV^ΔNS1^ or YFV^ΔCME^ required the expression of the complementation from the removed genes for trans-neuronal spreading, these viruses can directly infect the neurons at the injection site. Thus, while the current versions of YFV-17D are effective for tracing long-range neuronal projections, they may not be suitable for mapping local connectivity. (3) Complexity in applications. The genetically modified versions of YFV-17D necessitate the co-application of helper viruses, which complicates the overall procedure. (4) Capacity. Although it can carry a large gene, the 11kb genome of YFV-17D still sets an upper capacity limit of the cargo genes. In comparison, the HSV (including PRVs) can have larger cargos. (5) Biohazard materials. Despite YFV-17D being a well-characterized vaccine, it remains a live virus. Experimenters must be vigilant regarding potential biosafety issues and adhere to standard biosafety guidelines.

## Detailed Protocol

### Materials

#### ANIMALS

C57BL/6J mice from Jackson laboratory were housed in groups of 2–5 per cage, with each cage equipped with a nestlet and a mouse igloo for environmental enrichment. The housing room was maintained at a temperature of 23°C with a 12-hour light-dark cycle (lights on from 06:00 to 18:00). Food and water were provided *ad libitum*. ▲ CAUTION All animal experiments must be approved by the Institutional Animal Care and Use Committee (IACUC) at your institution and conducted in accordance with the *Guide for the Care and Use of Laboratory Animals* published by the National Research Council. At present, we have only tested the YFV-17D-derived viral vectors in mice. Given that YFV-17D originates from a human virus, these viral vectors are likely applicable to other mammalian models.

#### REAGENTS

##### Plasmids

###### Plasmids encoding YFV-17D genomic sequence

- pCCI-YFV^ΔNS1^-Cre (Addgene, Cat. No. 179952)
- pCCI-YFV^ΔNS1^-mVenus (Wei Xu lab, UT Southwestern)
- pCCI-YFV^ΔNS1^-mCherry (Wei Xu lab, UT Southwestern)
- pCCI-YFV^ΔCME^-mVenus (Addgene, Cat. No. 179953)
- pCCI-YFV^ΔCME^-mScarlet (Wei Xu lab, UT Southwestern)
- pCCI-YFV(P2A)^ΔCME^-mVenus (Wei Xu lab, UT Southwestern)
- pCCI-YFV(P2A)^ΔCME^-mScarlet (Wei Xu lab, UT Southwestern)
- pCCI-YFV^ΔCMENS1^-Cre (Wei Xu lab, UT Southwestern)
- pCCI-YFV^ΔCMENS1^-Flpo (Wei Xu lab, UT Southwestern)
- pCCI-YFV-mVenus (Charles Rice lab, Rockefeller University)
- pCCI-YFV-mScarlet (Wei Xu lab, UT Southwestern)

###### Plasmids for helper AAVs

- AAV-Syn-NS1 (Addgene, Cat. No. 175276)
- AAV-TRE-NS1NF (Addgene, Cat. No. 175279)
- AAV-TRE-NS1-dTomato (Addgene, Cat. No. 175278)
- AAV-TRE-C-prM-E-NS1 (Addgene, Cat. No. 175277)
- pAAV-Syn-rtTA (Addgene, Cat. No. 175274)
- AAV-Syn-tTA (Addgene, Cat. No. 175280)
- pAAV-DIO-NS1 (Addgene, Cat. No. 175273)
- AAV-Dlx-tTA (Addgene, Cat. No. 210730)

###### Plasmids for making packaging cell line

- FUW-C-prM-E-NS1: Addgene, Cat. No. 175281

▲ **Note** While we provide a detailed protocol for researchers to generate their viral vectors, the UCI Center for Neural Circuit Mapping Viral Core team (https://cncm.som.uci.edu/yellow-fever-virus) is ready to produce and distribute ready-to-use viral aliquots, including the YFV-17D-derived viruses and related helper AAVs to the broad neuroscience community on a recharge basis.

##### Reagents for amplification of pCCI plasmids

- TransforMax EPI300 Chemically Competent E. coli (Epicentre, Cat. No. C300C105, including SOC medium)
- CopyControl Induction Solution (Epicentre, Cat. No. CCIS125)
- Chloramphenicol (Sigma, Cat. No. C0378)
- Plasmid miniprep kit, (Qiagen, Cat. No. 4789)
- Plasmid Plus Midi Kit, (QIAGEN, Cat. No. 12945)

##### Reagents for in vitro transcription

- mMESSAGE mMACHINE™ SP6 Transcription Kit (Life Technologies, Cat. No. AM1340)
- Restriction endonuclease Xho1 and buffer (NEB, Cat. No. R0146L)
- Buffer-saturated phenol/chloroform (Sigma, Cat. No. P2069-400ML)
- Chloroform (Fisher, Cat. No. AC383770010)

##### Reagents for making BHK cells stably expressing NS1 or C-prM-E-NS1

- 293T cells (ATCC, Cat. No. CRL-3216)
- DMEM (Life Technologies, Cat. No. 11995065)
- Fetal bovine serum (FBS) (Life Tech, Cat. No. 26140079)

##### Reagents for transfection and collection of viral solution

- BHK cells (ATCC, Cat. No. CCL-10) or BHK cells stably expressing C-prM-E-NS1 (Wei Xu lab at UT Southwestern, or self-make with BHK cells and above lentiviral vectors FUW-C-prM-E-NS1)
- TransIT-mRNA Transfection Kit (Mirus Bio, Cat. No. MIR 2225)
- Amicon Ultra-15 Centrifugal Filter Unit 100 kDa NMWCO (Millipore, Cat. No. UFC910096)
- Opti-MEM (Life technologies, Cat. No. 31985062)
- DMEM (Life technologies, Cat. No. 11995065)
- PEG8000 (Calbiochem, Cat. No. 6520-5kg)

##### Reagents for qRT-PCR titering

- qRT-PCR Primers and probes: forward primer—GCCTCCCACATCCATTTAGT; reverse primer—CAGGTCAGCATCCACAGAATA; and probe -- CCGAACGCTGATTGGACAGGAGAA (with FAM 520 dye and ZEN / Iowa Black™ FQ quencher, custom order from IDTDNA)
- Applied Biosystems TaqMan Fast Virus One-Step Master Mix (Life Technologies, Cat. No. 4444432)

##### Reagents for surgery and animal care

- Sterile Saline Solution (Covetrus, Cat. no. C1880725)
- Povidone Iodine Surgical Scrub Solution (Dynarex)
- Ethyl Alcohol 200 Proof (Pharmco Products, Cat. No. 111000200CSPP)
- Mineral Oil (Sigma Aldrich, Cat. No. M3516)
- Mouse Buprenorphine SR (slow release) 0.5 mg/mL (UTSW Animal Resource Center)
- Diluted Mouse Carprofen 1.3 mg/mL (UTSW Animal Resource Center)
- Clorox Splash-Less Liquid Bleach, Regular Scent, 77 fl oz (Clorox)
- Isoflurane Inhalation Solution 99.9% (Piramal Critical Care, Cat. No. 66794001725)
- Custom research diet with doxycycline 625ppm (Teklad Envigo, Cat. No.TD.01306)
- Rapamycin (Fisher, Selleck Chemical, Cat. No. S1039)

#### EQUIPMENT

##### Procedure 1, Packing YFV-17D-derived viral vectors

▲ **CRITICAL** Most equipment may be substituted for equal alternatives.

###### Amplification of plasmid

- Incubated Shaker (Eppendorf, Cat. No. M1282-0000)
- Centrifuge (Thermo Scientific, Cat. No. 75818382)
- Falcon 14mL Round Bottom Polystyrene Test Tube (Life Sciences, Cat. No. 352057)
- 50ml Conical Centrifuge Tubes, Polypropylene Bulk, Sterile (Olympus Plastics, Cat. No. 28-108)
- Nanodrop 2000c Spectrophotometer (Thermo Scientific, Cat. No. ND2000)
- Desktop microcentrifuge (Thermo Scientific, Cat. No.75-772-436)

###### In vitro transcription

- Cell culture incubator (Panasonic, Cat. No. MCO-170AICUVL-PA)
- Class II biosafety cabinet (NUAIRE, Cat. No. NU-425-600)

###### Transfection and collection of viral solution

- Real-time PCR system (Applied Biosystems, Cat. No. A43054)

##### Procedure 2, Intracranial injection surgeries

▲ **CRITICAL** Most equipment may be substituted for equal alternatives.

###### Surgical equipment for Procedure 2

- Dual-Stage Glass Micropipette Puller PC-10 (Narishige, Cat. No. PC-10)
- 3.5” Nanoject Glass Capillaries (Drummond, Cat. No. 3-000-203-G/X)
- Nanoject 2” 30 Gauge needle (Drummond, Cat. No. 3-000-027)
- 1 mL BD Luer-Lok Syringe only (BD, Cat. No. 00382903096282)
- 3M Comply Lead-free Autoclave Steam Indicator Tape (3M Health Care)
- Parafilm M Sealing Film (Parafilm, HS234526B)
- Instant Sealing Sterilization Pouches (Fisherbrand, Cat. No. 01-812-50)
- Self-Sealing Sterilization Pouch 3.5” x 6.5” (RMH3 Dental)
- Kimberly-Clark Professional Kimtech Science Kimwipes Delicate Task Wipes (Kimtech)
- Standard Size Heating Pad (Sunbeam, Cat. No. 731-500-000R)
- Nexgen Mouse 500 (Nexgen, Cat. No. 223581-2)
- Stereotaxic “U” Frame Assembly (David Kopf, Model 900-U, with Digital Display)
- Leica M80 Stereomicroscope (Leica)
- Swingarm Stand (Leica)
- Nanoject III (Drummond, Cat. No. 3-000-207)
- Micro Bead Sterilizer with Glass Beads (Benchmark Scientific, Cat. No. B1201)
- Homeothermic Monitoring System for Small Rodents (Harvard Apparatus, Cat. No. 55-7020)
- Avante Compact 150 Rodent Anesthesia Machine (Avante, Cat. No. 10150)
- Revvity Health Sciences Inc XAF-8 Anesthesia System Filters (Fisher Scientific, Cat. No. NC0482435)
- Patterson Scientific Sliding Top Induction Chamber Small (Patterson Scientific, Cat. No. 78933389)
- USP Medical Grade Oxygen, Size E High Pressure Aluminum Medical Cylinder, CGA 970 Yoke Fitting (Airgas)
- Oil Only Sorbent Pads - 15 x 19”, Light (ULINE, Cat. No. S-12889)
- Kitchen Food Scale, LCD Digital Display, 8” x 6” x 1.25” (Amazon)
- 1 mL Allergy Syringe with Permanently Attached 27 G x 3/8 in. Intradermal Bevel Needle (BD, Cat. No. 00382903055418)
- World Precision Instrument Micro Mosquito Hemostatic Forceps, 12.5 cm, Curved, German (Fisher Scientific, Cat. No. 50-822-519)
- Straight Strong Point Tweezer 2-Star (Excelta, Cat. No. 00-PI)
- Micro-dissecting Scissors 4.25” (Sigma-Aldrich)
- 0.5 mm Dia Drill Burr (Fine Science Tools, Cat. No. NC9152343)
- Brush Type Control Unit, 110-120 Volt (Foredom, Cat. No. HP4-917)
- Cotton Tipped Applicator 6” Wood Handle Non Sterile (HSI, Cat. No. 1009175)
- MAJOR LubriFresh P.M. Lubricant Eye Ointment (MAJOR)
- Gel Cream Hair Remover (VEET)
- Sterile Alcohol Prep Pads (Dynarex)
- Research Plus Pipette (Eppendorf, Cat. No. 3123000020)
- 10 μL XL Non-Filtered Universal Pipette Tips (VWR)
- Applying / Removing Forceps 7,5mm Clips, 12,5cm (FST, Cat. No. 12029-12)
- Michael Suture Clips (FST, Cat. No. 12040-01)
- Surgical Loop 3-ply Face Masks (Concentric Health Alliance)
- Fisherbrand Powder Free Nitrile Gloves (Fisher Scientific, Cat. No. 19-130-1597A)
- Eye Protection Glasses/Shield
- Uline Isolation Gowns – Yellow (Uline, Cat. No. S-24019Y)

###### Cell culture

70% (vol/vol) ethyl alcohol. Dilute 100% ethyl alcohol with MilliQ water until 70% (vol/vol) solution is achieved. Solution can be stored at RT (20 - 25°C).

▲ **CAUTION** Flammable reagent. Store solution in approved safety containers

*Adherent BHK cells:* Prepare cell culture medium -- DMEM with 10% fetal bovine serum (FBS). Medium can be store at 4 °C for ≤1 month. Cell culture medium should be warmed to 37 °C before adding to cells. Thaw BHK cells according to suppliers’ instructions. Culture cells in T25 or T75 flasks and maintain cells at 37 °C with 5% CO2. Use 0.05% (vol/vol) trypsin-EDTA for trypsinization of cells during cell passaging. BHK cells should be transfected or passaged when 75%-90% confluent. Split cells 24 hours before transfection.

###### Plasmid amplification

LB medium: Mix 1% (wt/vol) NaCl, 1% (wt/vol) tryptone, and 0.5% (wt/vol) yeast extract in deionized water. Sterilize via an autoclave. Store for up to 1 year before use. Discard the medium if it becomes turbid.

LB agar plates: Mix 1.5% (wt/vol) agar, 1% (wt/vol) NaCl, 1% (wt/vol) tryptone, and 0.5% (wt/vol) yeast extract in deionized water. Sterilize via an autoclave. Cool to ∼50 °C and add chloramphenicol to achieve 15 µg/ml final concentration. Pour LB agar into 10-cm Petri dishes and store at 4 °C. The maximum recommended storage time for ampicillin-treated LB agar plates is 3 months.

###### Material setup for Procedure 2

Prepare sterile glass needles: Pull 3.5” Nanoject glass capillaries with Micropipette Puller (PC-10). Adjust the settings of temperature to achieve desired length and tip size of the injection needles. Fill the needles with mineral oil and autoclave for later surgical use. Autoclaved needles may be stored at RT (20 - 25°C) for up to 6 months unopened.

Prepare sterile parafilm: Cut Parafilm into small squares ∼15 mm x 15 mm. Place squares into sterilization pouch, making sure they do not overlap. Sterilize via gas autoclave sterilizer. Unopened autoclaved parafilm squares may be stored at RT (20 - 25°C) for 6 months.

Prepare 10% (vol/vol) Bleach for viral waste: Dilute an appropriate volume of bleach with water to achieve 10% (vol/vol) concentration. The diluted bleach remains effective within 24 hours.

Cotton-tipped applicator preparation: Sterilize cotton-tipped applicators before surgical use. Unopened autoclaved cotton-tipped applicators may be stored at RT (20 - 25°C) for 6 months.

###### Stereotaxic equipment setup

Assemble stereotaxic instruments according to manufacturer guidelines. Attach Nanoject III so that it is perpendicular to the surgery surface. Ensure all tubing is properly connected between oxygen, isoflurane, and absorption filter. Activated charcoal absorption filter should not be expired and have sufficient absorption capacity remaining. Prepare drill according to manufacturer guidelines.

#### PROCEDURE

##### Procedure 1: Packaging of viruses (Fig. 3)

▲ **CAUTION** YFV-17D is a live attenuated vaccine. Both YFV-17D and the viral vectors engineered from it are classified as “Risk Group 2 agents” under the *NIH Guidelines for Research Involving Recombinant or Synthetic Nucleic Acid Molecules* (NIH Guidelines). The *Biosafety in Microbiological and Biomedical Laboratories* (BMBL), 6th edition, offers comprehensive guidance on handling Risk Group 2 pathogens, including laboratory practices, transportation, and procedures for associated animal studies.

###### Day 1: Transformation of pCCI plasmids

1 Thaw TransforMax EPI300 cells competent cells on ice.
2 Pre-chill a 14 ml Falcon tube on ice.
3 Add 25-50 ng of DNA and 25 µl of competent cells to the bottom of each of the pre-chilled tube.
4 Incubate on ice for 30 minutes.
5 Heat shock for 30 seconds at 42° C. ▲ **CRITICAL** The duration and temperature of the heat shock must be precise, as deviations—such as extended duration or altered temperature—can significantly reduce transformation efficiency.
6 Incubate on ice for 1 minute.
7 Add 200 ml SOC medium to the tube.
8 Shake at 37°C for 60 minutes. ▲ **CRITICAL** This recovery step is essential and should neither be omitted nor shortened.
9 Spread 50 µL of the cells onto an LB agar plate with 15 µg/ml of chloramphenicol.
10 Incubate the plate overnight (15-16 hours) at 37° C. ◆ **TROUBLESHOOTING**

###### Day 2: Inoculation

11 Inoculate a single colony into 10 ml of LB medium with 15 µg/ml of chloramphenicol.
12 Incubate the culture at 37°C with shaking at 220–250 rpm for 12–16 hours

###### Day 3: Copy control induction and midi-prep

13 Combine 10 ml overnight culture and 90 ml of LB media with 15 µg/ml of chloramphenicol, and 100 µl of CopyControl Induction Solution in a 500ml glass flask.
14 Grow the bacteria at 37°C for 5-6 hours with vigorously shaking (225-250 rpm). ▲ **CRITICAL** Shortening or extending the 5–6-hour incubation time may reduce DNA yield.
15 Collect the bacteria by centrifuging at 3000 × g for 45 minutes, then extract the plasmid using a DNA purification midi kit following the low-copy, large-plasmid protocol. Typically, 50 mL of bacterial culture is sufficient for one midi-prep.
16 Measure the DNA concentration with Nanodrop. Normally 100-300 µg of DNA can be purified with the bacteria from 50 ml of bacteria culture. ◆ **TROUBLESHOOTING**

###### In vitro transcription

17 Linearize the plasmid with Xho1. Set up the following reaction:

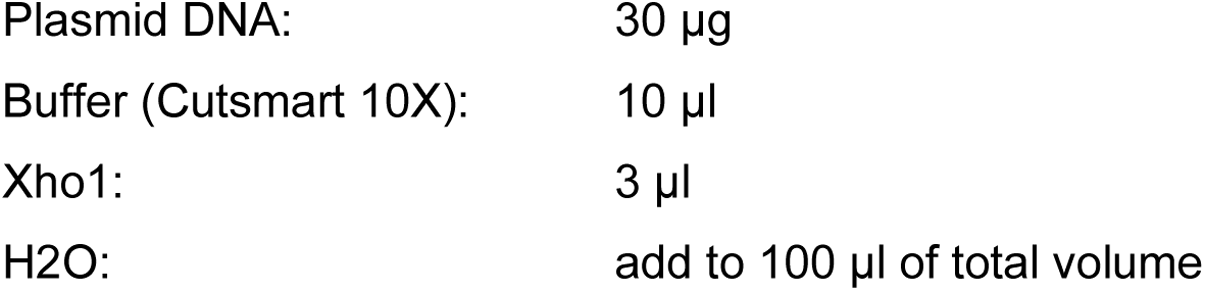 Incubate at 37 ⁰C for 1 hour. Terminate the restriction digest by adding 5 µl of 0.5 M EDTA, 10 µl of 3 M sodium acetate and 200 µl of ethanol. Mix well.
18 Precipitate the DNA by placing the above mixture at –20°C for at least 30 min. Then pellet the DNA in a desktop microcentrifuge at top speed (14,000RPM) for 15 min. Remove the supernatant, re-spin the tube for a few seconds, and remove the residual fluid with a fine-tipped pipet tip. Air dry the DNA for 10-15 min and then re-suspend the DNA in 30 µl of H2O. Measure the concentration of plasmids. Expected yield: normally 15-25 ug of DNA can be recovered.
19 In vitro transcription (with mMESSAGE mMACHINE® Kit) In RNase-free conditions, add the following components sequentially:

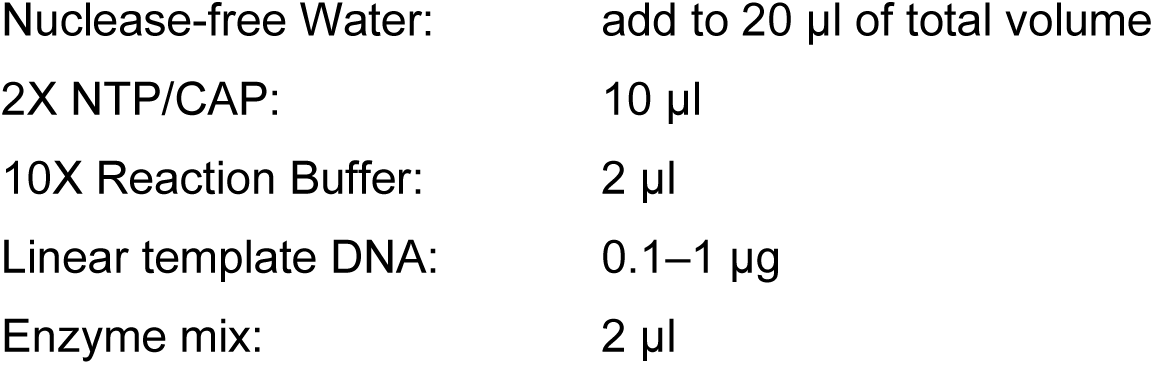 Incubate at 37 ⁰C for 2 hours.
20 Add 1 μl of TURBO DNase, mix well and incubate at 37°C for 15 min.
21 Add 115 μl of nuclease-free water and 15 μl of ammonium acetate stop solution (provided in the kit), and mix thoroughly.
22 Add 150 μl of buffer-saturated phenol/chloroform, and 150 μl of chloroform, mix by vortex vigorously. Centrifuge in a microcentrifuge at top speed (14,000RPM) for 5 min. Transfer the aqueous phase (upper phase) to a new tube.
23 Precipitate the RNA by adding 150 μl of isopropanol. Chill the mixture at –20°C for at least 15 min. Centrifuge at 4°C for 15 min at maximum speed (14,000RPM) to pellet the RNA. Carefully remove the supernatant and resuspend the RNA in 20 µl of nuclease-free water.
24 Measure the concentration of RNA and store the RNA at –80°C. Expected yield: 5-15 ug of RNA/reaction. The reaction may be duplicated to ensure an adequate yield. In case a large preparation of YFV is needed, the in vitro transcription reaction can be scaled up accordingly. ▲ **CRITICAL** Take care to avoid contamination with RNase in Steps 17-24. ◆ **TROUBLESHOOTING**

###### Option 1, small-scale preparation

25 Grow and maintain BHK cells in DMEM with 10% FBS.
26 Split confluency BHK cells 1:9 (by culture areas, the cells will be 30-40% confluency at the time of transfection) to 6-well culture plates the day before transfection.
27 Prepare transfection reagents (with TransIT-mRNA Transfection Kit; the amounts are for each well in the 6-well plate, with 2 ml of medium in the well):

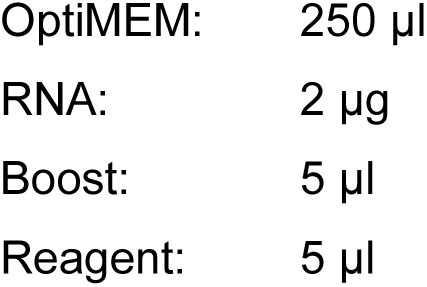 Incubate at room temperature for 3 min.
28 Add the above mixture drop-wise into the wells. Return to cell culture incubator.
29 Change to fresh cell culture medium one day. ▲ **CRITICAL** If the medium is not changed, the viral RNA used for transfection may be detected during titration by qRT-PCR, leading to an overestimation of the titer.
30 Collect the cell culture media from each of the wells 72 hours after transfection (Fig. 4).
31 Centrifuge the collected cell culture media at 3,000g for 30 min and transfer the supernatant to a 15 ml 100 KD MWCO filter. Centrifuge the filter at 3,000g for 3-10 min and concentrate the solution to ∼300 µl. Aliquot the solution containing YFV-17D vectors (10 µl/tube) and store at −80°C ▲ **Caution** The centrifugation time for the MECO filter may vary depending on the viral preparation. To avoid over concentration of the solution, centrifuge in 1- to 2-minute intervals.

###### Option 2, large-scale preparation

32 Grow and maintain BHK cells in DMEM with 10% FBS.
33 Split BHK cells 1:9 to T225 flasks 16 hours before transfection (the cells will be 30-40% confluency at the time of transfection).
34 Prepare transfection reagents (with TransIT-mRNA Transfection Kit; the amounts are for each T225 flask):

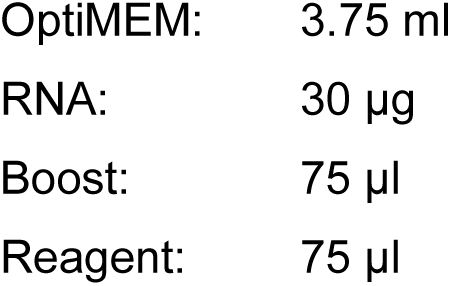 Incubate at room temperature for 3 min.
35 Add the above mixture drop-wise into T225 flasks. Return the flask to cell culture incubator.
36 Change to fresh cell culture medium one day after transfection. ▲ **CRITICAL** If the medium is not changed, the viral RNA used for transfection may be detected during titration by qRT-PCR, leading to an overestimation of the titer.
37 Collect the cell culture media from each of the T225 flasks into a 50-ml centrifuge tube 72 hours after transfection.
38 Prepare 40% PEG solution as the following:

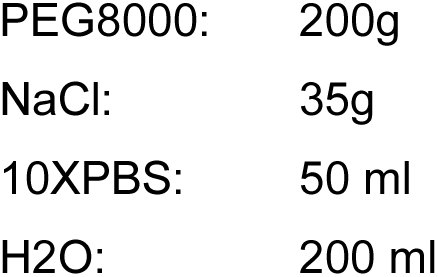 Mix by stirring at 50 °C until the chemicals are fully dissolved, adjust the pH to 7.20, and bring the final volume to 500 ml with H_2_O. Sterilize by running the solution through 0.2 µm filter. ▲ **Caution** It takes a few hours to prepare 40% PEG solution. The solution can be prepared ahead of time and stored at 4°C for 2-3 months.
39 Centrifuge the collected cell culture media at 3,000g for 30 min and transfer the supernatant (30 ml) to a new 50-ml centrifuge tube. Mixed with 7.5 ml of 40% PEG (medium: 40%PEG = 4:1 by volume) and kept on ice with rotation for >10 hours.
40 Centrifuge at 3500g for 60 min. Removed the supernatant containing PEG. Wash the pellet by carefully overlaying 150 µl of OptiMEM on top of the pellet and then pipette all solution out.
41 Resuspend the pellet by adding 1.0 −1.5 ml of OptiMEM, keep the tubes on ice for 1 hour, shake briefly each 5-10 min. Gently pipetting the mix with 1-ml pipette for 10-20 times to promote the dissolving of the pellet. In case a significant portion of the pellet remains undissolved, add more OptiMEM.
42 Centrifuge at 3000g for 30 min to remove undissolved materials in the mix. Aliquot the cleared supernatant solution containing YFV-17D vectors (10 µl/tube) and store at −80°C. ◆ **TROUBLESHOOTING**

###### Step 4: Titration of viruses

43 Prepare DNA standards: Linearize the pCCI plasmid with Xho1 as described in Step 2. Measure the concentration and serially dilute to 10 ng/µl, 1 ng/µl, 100 pg/µl, 10 pg/µl, 1 pg/µl, 100 fg/µl and 10 fg/µl of standards. Calculate the copy number based on molecular weight and Avogadro constant.
44 Dilute the YFV-17D solution by 1,000 fold and incubate at 95°C for 10 min.
45 Load 1 µl of viral sample or DNA standard, in duplicate or triplicates. Per 20 µl reaction:

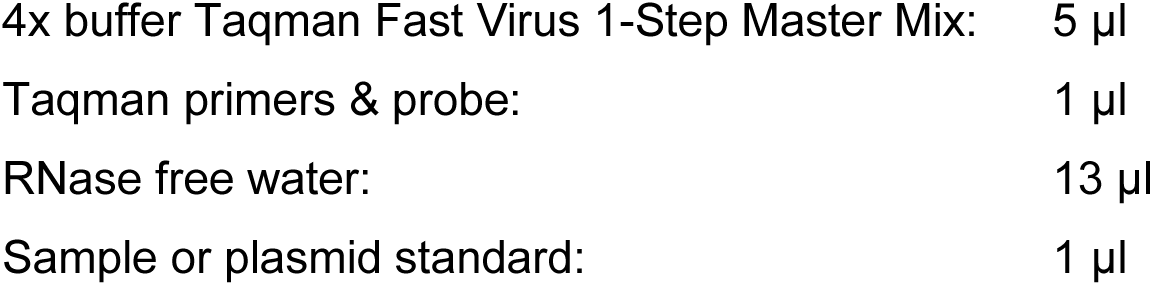
46 Run AQ (Absolute quantification) qRT-PCR in ABI 7500 Fast System with the following thermal cycles:

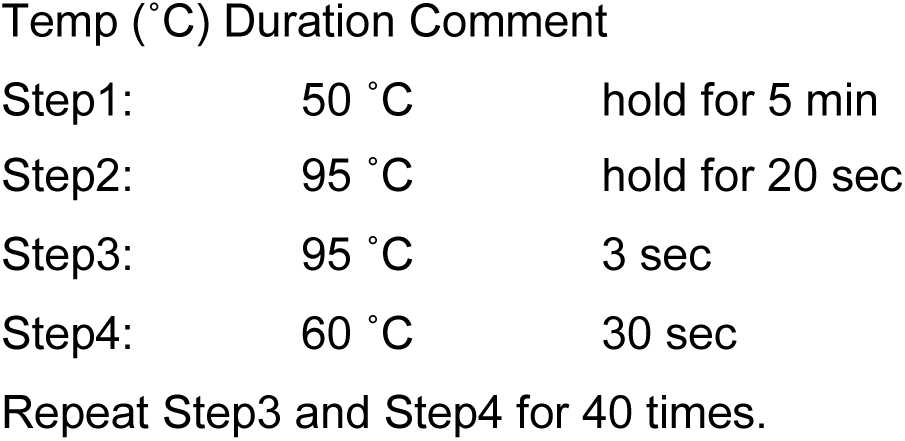
47 Run AQ analysis and record the titer. Expected yield: 1.0 X 10^10^ to 5.0 X 10^11^ genomic copies per ml. ▲ **Caution** The infectious titer of some of the viruses can also be measured by infection of BHK cells with serial diluted viral solution. The infectious titers are normally 1000 to 10,000-fold smaller than the genomic titers measured with qRT-PCR.

##### Procedure 2: In vivo application of YFV-17D-derived vectors

▲ **CAUTION** The procedures for *in vivo* application of YFV-17D-derived vectors vary based on the experimental objectives and the specific version of the viral vector chosen (Fig. 5). Many experiments require two sequential surgeries. The interval between the two surgeries is a critical factor influencing the efficiency of trans-neuronal spreading. In cases where the interval cannot be adjusted (e.g., when additional time is needed to activate genes carried by helper AAVs), administering medications such as rapamycin to reduce antiviral immune responses around the time of injection of YFV-17D-derived vector can enhance trans-neuronal spreading efficiency (Fig. 5D).

###### TIMING 10 – 15 min

▲ **CRITICAL** Mice should be housed according to institutional standards, and surgical tools should be sterilized before use.

▲ **CAUTION** All experiments using animals must be approved by IACUC and performed based on the guidelines set by the government and institutions.

▲ **CAUTION** Surgery protocol involves the use of AAV and YFV-17D-derived viruses. All procedures must comply with guidelines and regulations set by the Institutional Biosafety Committee for biosafety.

48 Turn on stereotaxic surgery equipment and bead sterilizer. Prepare a 10% bleach solution. Set the heating pad (placed in the stereotaxic frame, will be underneath the mouse) to 37°C and place a clean recovery cage without any bedding. The recovery cage is placed partially on top of a heating pad (setting at 37°C) so that the mouse can move between warmer and colder areas in the cage.
49 Weigh the mouse and prepare carprofen (5 mg/kg) and buprenorphine SR (1.0 mg/kg) based on the body weight.
50 Place the mouse in induction chamber and induce anesthesia with 3.5-4% (wt/vol) isoflurane.
51 When the mouse stops voluntary movements, transfer the mouse to stereotaxic frame with continuous inhalation of 1.0-2.0% of isoflurane, regularly monitoring the breathing rate and adjusting the dosage of isoflurane to maintain sufficient anesthesia and avoid overdose. Apply artificial eye lubricant to the animal’s eyes. **▲ CRITICAL** Confirm the mice are fully anesthetized with toe-pinch reflex test. ▲ **CRITICAL** The animal’s head should be properly aligned on the stereotactic frame, as this is essential for accurate injections.
52 Apply depilatory cream on the skull over a surface at least 1.5 times bigger than the surgery area and wait for 45 seconds before removing with a rubbing alcohol pad and gauze. **▲ CRITICAL** Depilatory cream should not be left on animal skin for over 45 seconds as the risk for chemical burns increases. Ensure the removal of all depilatory cream before proceeding.
53 Disinfect the surgical site with alternating applications of betadine and 70% ethanol at least three times.
54 Administer carprofen and buprenorphine SR subcutaneously. ◆ **TROUBLESHOOTING**

###### TIMING 15 – 30 min

55 Confirm the animal is fully anesthetized by toe pinch test and adjust isoflurane appropriately.
56 Make a midline incision with fine scissors and tweezers and expose the skull to identify bregma and lambda. Clean off any excess connective tissue on the skull with cotton tips for better visualization. Adjust the ear bars and use the midline of the skull (the Bregma-Lambda connection line) to assure mouse skull is properly aligned on the stereotactic frame to ensure accurate stereotaxic coordinates.
57 Adjust the stereotaxic nose cone to align the Bregma and Lambda on the same horizontal plane, ensuring a vertical discrepancy of no more than 0.1 mm.
58 Drill a burr hole (∼ 1-2 mm^2^) on the skull in the region on top of the targeted brain regions to expose the dura. **▲ CRITICAL** Drilling should be careful to avoid any damage of dura or the brain tissue.
59 Install glass capillary filled with mineral oil onto the nanoinjector. Place a small piece of sterilized parafilm on the surface of the mouse head and pipette an appropriate amount of viral solution (0.75-1.25 µl) onto the parafilm. The virus should be placed on ice until it’s ready to be used. Load proper volume of viral solution (0.5-1.0 µl) from the parafilm into the glass capillary. Ensure there is no air in the capillary.
60 Position the needle at the appropriate anterior-posterior and medial-lateral coordinates relative to Bregma. Use the Allen Brain Reference Atlas (2008 version) or the Paxinos Mouse Brain Atlas to determine the coordinates.
61 Lower the needle until it just touches the surface of the dura and use this as the zero point for the dorsal-ventral axis. Slowly advance the needle to penetrate the dura and reach the targeted depth. And then inject the viral solution at a rate of 1 nL/s, with the injection volume ranging from 0.25 to 0.75 µL, depending on the size of the target region and the experimental objectives.
62 Leave the needle in place for at least 5 minutes to allow the virus to diffuse into brain tissue.
63 Slowly withdraw needle from the brain. Once completely withdrawn, discard the injection needle. Caution: All materials that contacted viral solutions, such as pipette tips, glass needles, and parafilm, must be decontaminated in a 10% bleach solution before disposal.
64 Close the incision of the skin using sterile surgical staples or suture.
65 Place the mouse in the recovery cage prepared earlier and wait until it fully wakes up and can voluntarily move around inside the cage. It normally takes 10-30 min.
66 For certain version so the viruses, the mouse should be placed on a Dox diet as specified in Fig. 5.
67 Follow institutional guidelines for post-surgical documentation and clean workspace. ◆ **TROUBLESHOOTING**

###### TIMING 1 – 2 weeks

68 Monitor the mouse for signs of distress or infection following at 24, 48 and 72 hours after injection and provide pain relief as needed.
69 24 hours later, provide pain relief with a subcutaneous injection of carprofen (5 mg/kg).
70 If surgical staples are used to close the skin incision, these staples need to be removed after 7 - 10 days.
71 Provide doxycycline diet if needed (as specified in Fig. 5).
72 Provide rapamycin (6mg/kg, Intraperitoneal injection, once every 24 hours for a total of 7 consecutive days) if needed (as specified in Fig. 5).
73 Wait for 8-12 days before subjecting the mice to histological analysis or functional studies of brain circuits. Mice designated for histological analysis are euthanized with 5% isoflurane, followed by intracardiac perfusion with 10 mL of PBS and 40 mL of 4% paraformaldehyde.

###### TIMING 25 – 35 min

74 Following 3-14 days after the initial surgery (the intervals between injections vary based on the viral version as specified in Fig. 5), repeat the steps 48-67 for the second surgery of stereotaxic injection, and the steps 68-73 for postsurgical recovery, histological analysis or other experimental procedures. ◆ **TROUBLESHOOTING**

## ANTICIPATED RESULTS

The YFV-17D-derived viral vectors are powerful tools for two major areas of neuroscience research: neuroanatomical mapping of brain circuits and targeting postsynaptic neurons for functional studies in living animals. The vectors (either replication-capable, replication-deficient, or packaging-deficient versions) carrying fluorescent proteins enable anatomical analysis of neuronal circuits. Following successful delivery to the region of interest, fluorescent labeling of neurons is observed both at the injection site and in downstream brain regions. The fluorescent signals are robust enough to reveal fine morphological details of labeled neurons without requiring immunohistochemical amplification (Fig. 6). In addition, YFV^ΔNS1^-Cre is a valuable tool for tracing astrocytes associated with specific neuronal pathways. When used in fluorescent reporter mouse lines, Cre recombinase activity induces strong expression of fluorescent proteins, facilitating the visualization of traced astrocytes. However, reporter AAVs have shown limited efficiency in reporting Cre activity from YFV^ΔNS1^-Cre in postsynaptic astrocytes. Most YFV-17D vectors have been successfully tested in various brain regions, including the cortex, hippocampus, thalamus, hypothalamus, midbrain, and brainstem structures. However, they exhibit limited efficiency in the cerebellum.

For functional analysis of brain circuits, YFV^ΔNS1^-Cre and YFV^ΔCMENS1^-Cre (or their Flpo counterparts) are the preferred viral vectors. These vectors spread to postsynaptic neurons and turn on (or turn off) Cre- or Flpo-dependent genes. The activated genes may encode sensors for monitoring neuronal activity, such as calcium indicator GCaMP, or effectors for modulating neuronal activity or other biological features, such as channelrhodopsins. Many of these sensors and effectors are fluorescently tagged, allowing straightforward confirmation of their activation by YFV^ΔNS1^-Cre and YFV^ΔCMENS1^-Cre. To minimize potential neuronal toxicity, the replication of YFV^ΔNS1^ can be stopped using doxycycline (Dox) after the desired genes have been turned on or off. Since YFV-17D is an RNA virus, its genomic RNA degrades rapidly within host cells. Typically, the viral RNA becomes undetectable in the brain within two weeks after replication is stopped. At this point, the mice become virus-free, while the activated reporter genes remain functional, enabling sustained monitoring or control of neuronal activity.

We encountered two major situations contributing to low labeling efficiency or high variability across experiments. 1. Inaccurate targeting during surgeries: For replication- or packaging-deficient versions of the viral vectors, two sequential surgeries are required. Precise targeting of both injections to the same brain locus is essential for efficient trans-neuronal spreading. Deviations in injection accuracy can significantly impact labeling outcomes. 2. Suboptimal timing of the experimental procedure: The temporal interval between injections plays a critical role in the efficiency of trans-neuronal spreading (Fig. 5). For instance, when using YFV^ΔCME^-mScarlet in combination with AAV-TRE-CMENS1 and AAV-Syn-tTA to map monosynaptic projections, a 3-day interval between the two injections was found to yield optimal results. Shorter or longer intervals led to reduced trans-neuronal spreading efficiency. Addressing these factors is crucial for achieving consistent and reliable experimental outcomes.

**Table 1.**
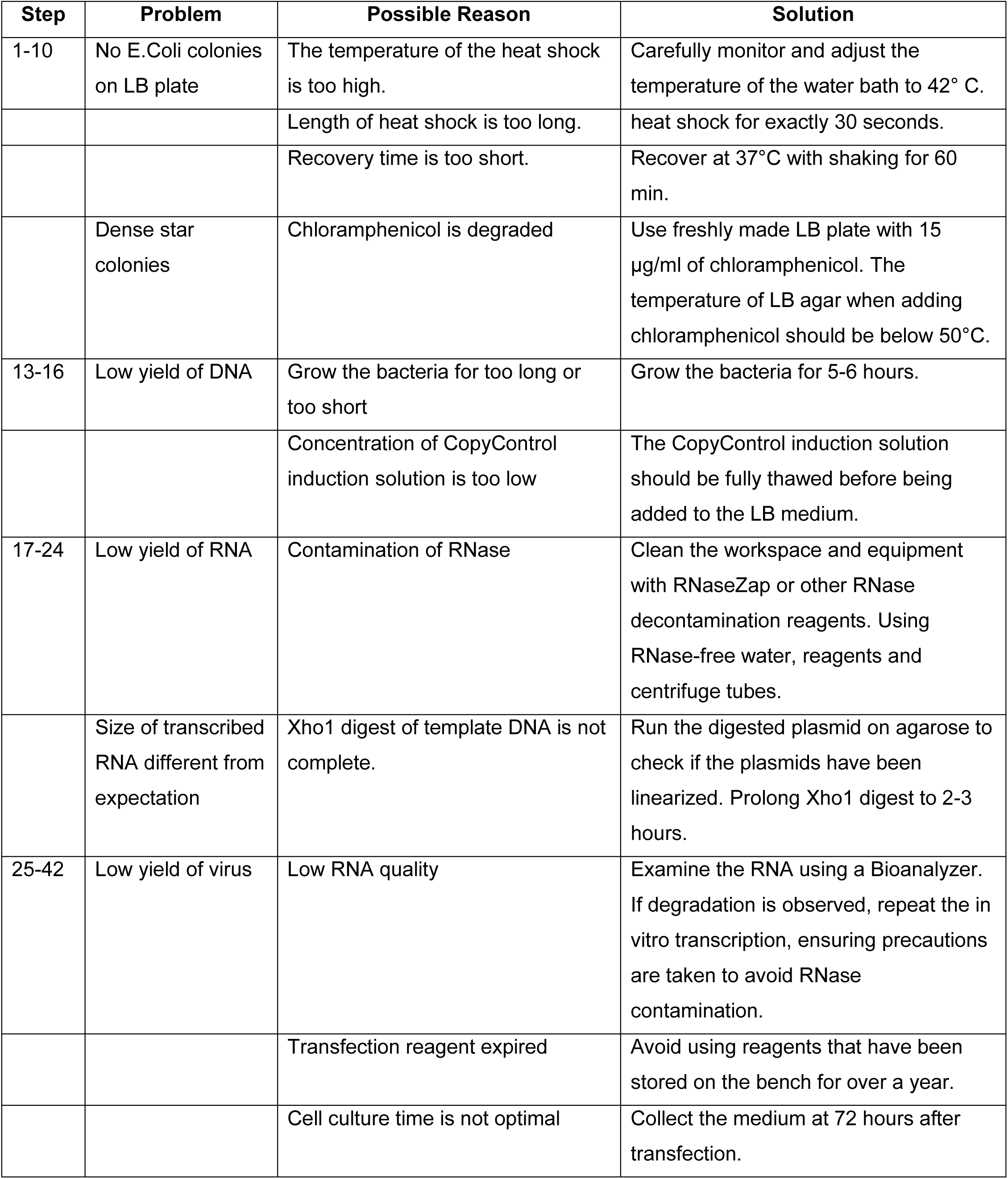
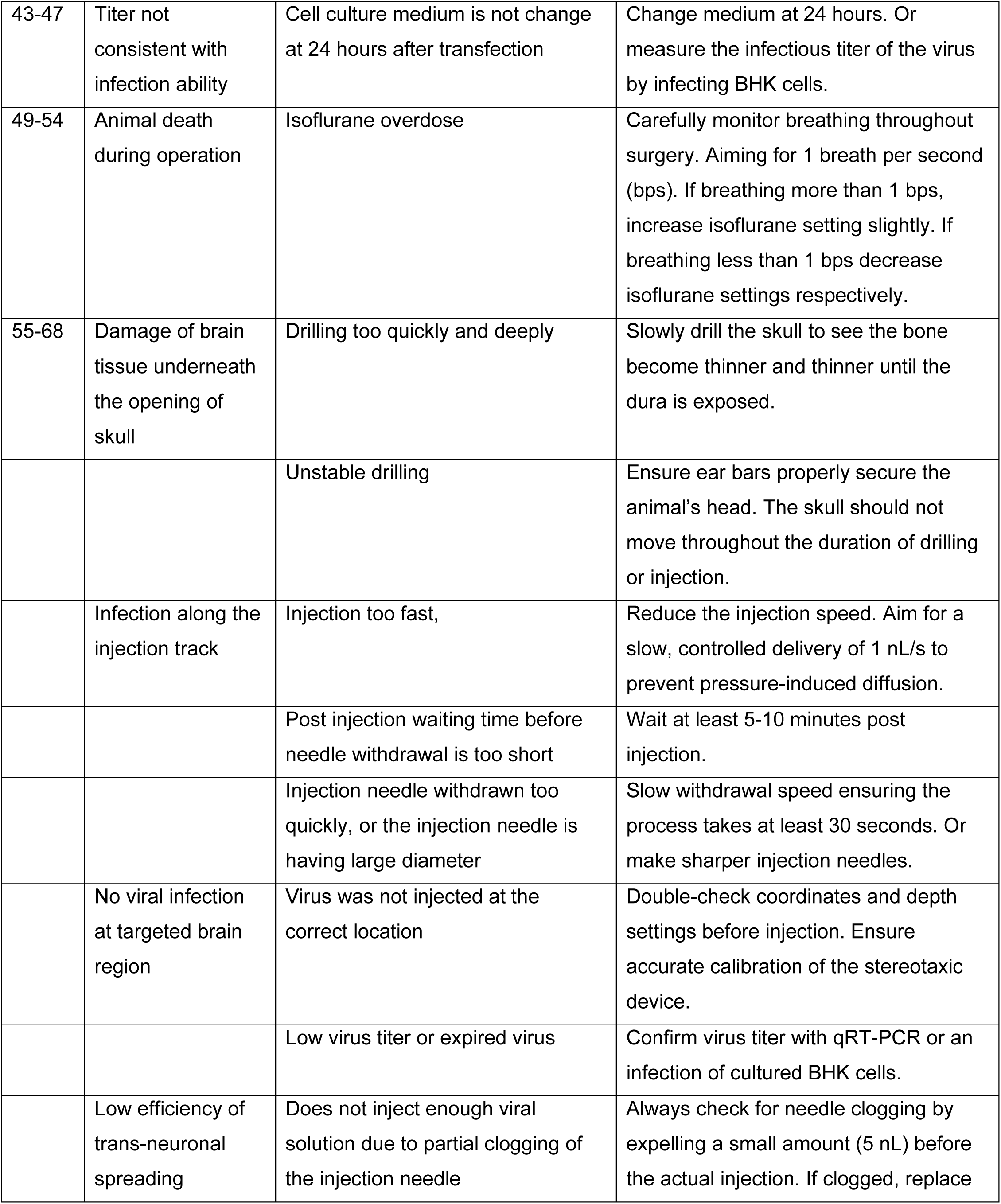
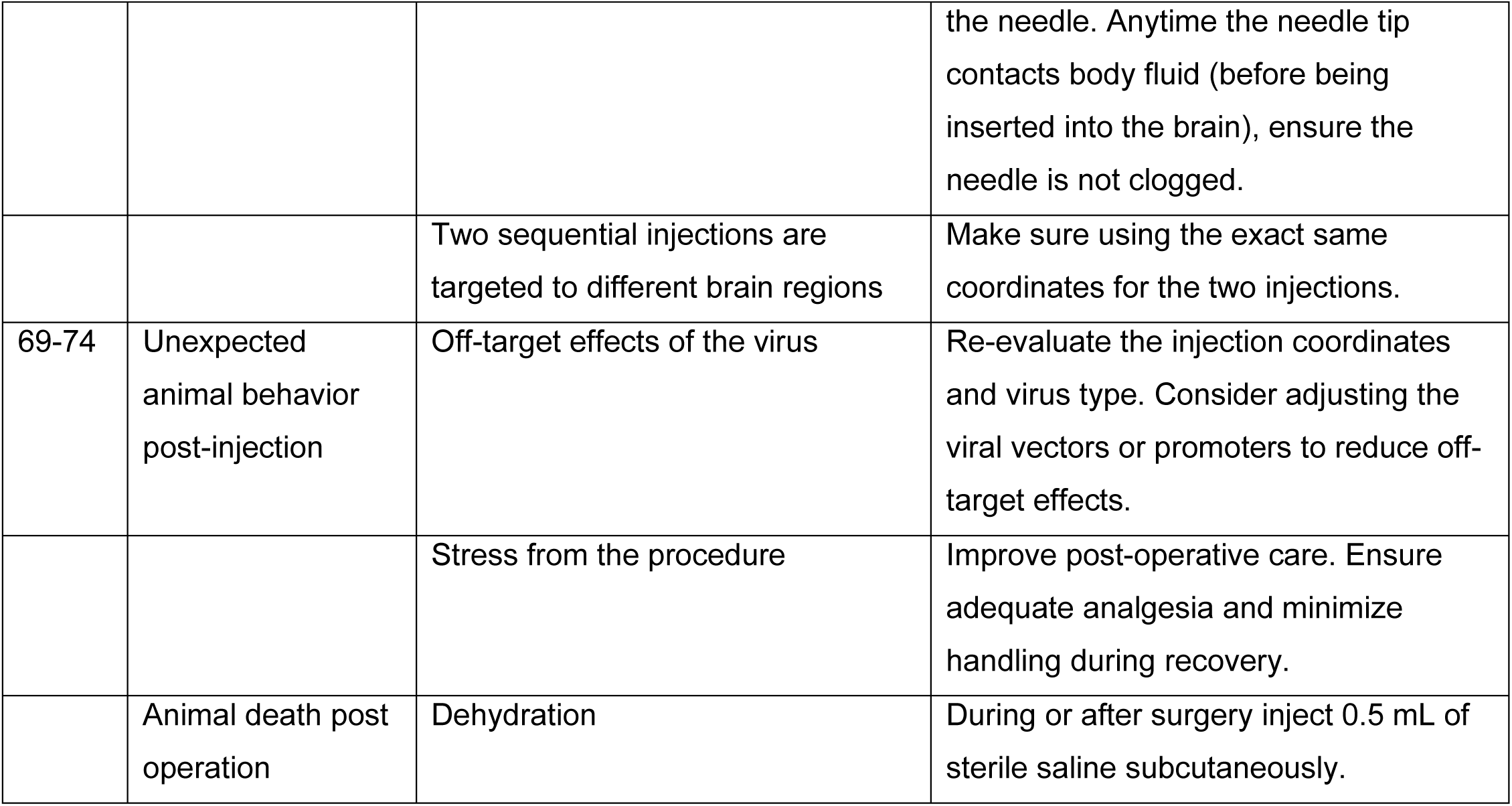
Troubleshooting table.

## Ethics approval and consent to participate

All animal work described in this study Animal work was approved and conducted under the oversight of the UT Southwestern Institutional Animal Care and Use Committee (IACUC) and complied with Guide for the Care and Use of Laboratory Animals by National Research Council.

## Consent for publication

all authors have approved the manuscript for submission.

## Availability of data and material

All data and materials described in this study are available upon request from the corresponding author.

## Author contributions

WX designed and supervised the study. TP, LM, US, AL, IN, YL, and WX carried out the research. All authors contributed to the manuscript preparation.

## Acknowledgements

This project was supported by a grant from NIH (1RF1MH130422 and 1U01AG076791). We thank Dr. Denise Ramirez (Whole Brain Microscopy Facility, UT Southwestern) and Dr. Shin Yamazaki (Neuroscience Microscopy Facility, UT Southwestern) for their assistance with brain section imaging. We also extend our gratitude to the broader neuroscience community, especially the researchers at the UCI Center for Neural Circuit Mapping, for their insightful discussions and feedback.

